# Sub-second multi-channel magnetic control of select neural circuits in behaving flies

**DOI:** 10.1101/2021.03.15.435264

**Authors:** Charles Sebesta, Daniel Torres, Boshuo Wang, Joseph Asfouri, Zhongxi Li, Guillaume Duret, Kaiyi Jiang, Zhen Xiao, Linlin Zhang, Qingbo Zhang, Vicki Colvin, Stefan M Goetz, Angel V Peterchev, Herman Dierick, Gang Bao, Jacob T. Robinson

**Affiliations:** Department of Bioengineering, Rice University, Houston, Texas, USA; Department of Electrical and Computer Engineering, Rice University, Houston, Texas, USA; Department of Psychiatry & Behavioral Sciences, School of Medicine, Duke University, Durham, North Carolina, USA; Department of Electrical and Computer Engineering, School of Engineering, Duke University, Durham, North Carolina, USA; Department of Neurosurgery, School of Medicine, Duke University, Durham, North Carolina, USA; Department of Biomedical Engineering, School of Engineering, Duke University, Durham, North Carolina, USA; Department of Neuroscience, Baylor College of Medicine, Houston, Texas, USA; Department of Human and Molecular Genetics, Baylor College of Medicine, Houston, Texas, USA; Department of Chemistry, Brown University, Providence, Rhode Island, USA

## Abstract

Precisely timed activation of genetically targeted cells is a powerful tool for studying neural circuits and controlling cell-based therapies. Magnetic control of cell activity or “magnetogenetics” using magnetic nanoparticle heating of temperature-sensitive ion channels enables remote, non-invasive activation of neurons for deep-tissue applications and studies of freely behaving animals. However, the *in vivo* response time of thermal magnetogenetics is currently tens of seconds, which prevents the precise temporal modulation of neural activity similar to light-based optogenetics. Moreover, magnetogenetics has not provided a means to selectively activate multiple channels to drive behavior. Here we produce sub-second behavioral responses in *Drosophila melanogaster* by combining magnetic nanoparticles with a rate-sensitive thermoreceptor (TRPA1-A). Furthermore, by tuning the properties of magnetic nanoparticles to respond to different magnetic field strengths and frequencies, we can achieve sub-second, multichannel stimulation, analogous to multi-color optogenetic stimulation. These results bring magnetogenetics closer to the temporal resolution and multiplexed stimulation possible with optogenetics while maintaining the minimal invasiveness and deep-tissue stimulation only possible by magnetic control.

## Main

Magnetic stimulation of genetically targeted cells, or “magnetogenetics”, may enable researchers to apply a magnetic stimulus throughout the brain of a freely moving animal in a non-invasive manner to study circuits that are deep within the brain or distributed over large areas. One approach of magnetogenetics with well described physical phenomena relies on two components to be present in the tissue: synthetic magnetic nanoparticles that convert alternating magnetic fields into heat and thermoreceptors that convert the local heat into neural activity^1–3^. While there have been reports of magnetogenetic technologies that rely on purely genetically encoded proteins^4–7^, it is currently unclear how these magnetically sensitized chimeric proteins function^8,9^.

Compared to optical methods for stimulating genetically targeted cells (optogenetics)^10,11^, magnetogenetics offers unique advantages for deep volumetric targets. While optogenetics has response times of 10-20 ms^12^, most optical wavelengths are only effective at distances of a few mm from an optical source due to tissue scattering. In contrast, magnetic fields in the frequency range of 0.1 – 1 MHz have very low attenuation in bone, air, and biological tissue^13^. This superior bone and tissue penetration of magnetic fields eliminates the need for invasive surgeries to introduce light probes typically required for optogenetic stimulation, interventions that can cause potential tissue damage from implantation and heat generation.

While magnetogenetics offers advantages including deep tissue volumetric stimulation and minimal invasiveness, the reported *in vivo* response time of magnetogenetic technologies is on the order of ten seconds – more than 1000-fold slower than optogenetic stimulation largely due to the thermoreceptors used. Previous experiments with membrane targeted cobalt-doped nanoparticles have shown latencies of 2.18 ± 0.17 s in *trpV1^+^* neurons *in vitro* and a 22.8 ± 2.6 s latency *in vivo* via motor cortex stimulation resulting in an ambulatory response in *trpV1^+^* mice^2^. Earlier experiments with undoped iron oxide nanoparticles showed a ~5 s latency *in vitro* with *trpV1^+^* neurons with upregulation of c-Fos expression *in vivo* on the order of minutes^3^. Existing magnetogenetic methods rely on thermoreceptors (e.g. TRPV1) that respond at temperatures several degrees above body temperature but heating the surrounding tissue to the threshold response temperature can take several seconds. These multi-second latencies prevent precise timing with behavioral or environmental cues that are essential for studying the relationship between neural activity and behavior. Magnetic activation of mechanoreceptors in contact with magnetic particles that move in response to a magnetic field offers a path to faster stimulation^14^, but the *in vivo* response time remains on the order of several seconds and requires micron-sized particles or aggregates that can be difficult to deliver *in vivo*^15^.

In this work we replaced threshold thermoreceptors with a rate-sensitive thermoreceptor to achieve sub-second response times approaching what can be achieved with optogenetics. Since magnetic nanoparticle heating can increase the tissue temperature rapidly, using thermal rate sensors eliminates the wait-time required for the tissue to reach a threshold activation temperature when using thermoreceptors like TRPV1. Recent work demonstrates that *Drosophila* TRPA1-A is activated by subtle temperature changes for temperature avoidance^16^. Additionally, this activation is susceptible to the rate of temperature change and rapid heating can lower the response threshold from ~34.5 °C to ~29.1 °C potentially because of calcium driven TRPA1 inactivation^17^. Additional experiments suggest that when natively expressed in organs or tissues^18^ TRPA1 is responsible for diverse sensitivity to temperature^19^ with behavioral responses to changes of 0.01 °C in *Drosophila^20^,* 0.005 °C in *C. elegans^21^,* and 0.003 °C in snakes^22^, making it an ideal target receptor to confer rapid, sensitive thermosensation. Therefore, we selected TRPA1-A as the thermoreceptor to optimize magnetothermal channel activation and demonstrate sub-second multi-channel magnetogenetics in *Drosophila.* Since TRPA1-A is native, rate sensitive, and commonly used for thermogenetics with genetic lines readily available, our approach can be applied to a wide range of magnetogenetics studies of brain function. *Drosophila* are used here to develop new magnetogenetic tools which can be adapted to other organisms. In larger animals, local heating of nanoparticles associated to rate sensitive thermoreceptors can stimulate targeted cells without the surrounding tissues being affected by bulk heating. For this test bed, we have chosen to modulate two easily observable phenotypes by activating cells expressing TRPA1 under the control of different *drivers:* 1) *Fruitless –* resulting in wing extension and 2) *Hb-9 –* resulting in side-to-side movement.

To test if the *Drosophila* TRPA1-A rate-sensitive thermoreceptor would indeed enable sub-second magnetogenetic activation we developed a system to measure *Drosophila* behavior under the influence of an alternating magnetic field (AMF). We generated fly strains that express thermal rate sensitive TRPA1-A channels under the control of the *fruitless* driver, which is known to control courtship behavior in males. Activation of cells expressing *fruitless* can be easily observed by a lateral wing extension behavioral response, as was previously shown with optogenetic^23,24^ and thermogenetic^25^ stimulation. Moreover, the behavior can be automatically tracked using pose estimation tools like DeepLabCut^26^ or FlyTracker^27^ eliminating observer bias. Instead of externally heating flies to activate the thermosensitive channel, we injected nanoparticles suspended into artificial *Drosophila* hemolymph (**Fig. 1A**, **S1**, see Methods). We placed the injected flies over an induction coil (**Fig. 1B**) and monitored the wing-opening behavior during AMF stimulation. The dissipated heat generated by the stimulated nanoparticles activates the dTRPA1-A protein channel (**Fig. 1C, D**).

**Figure 1.**
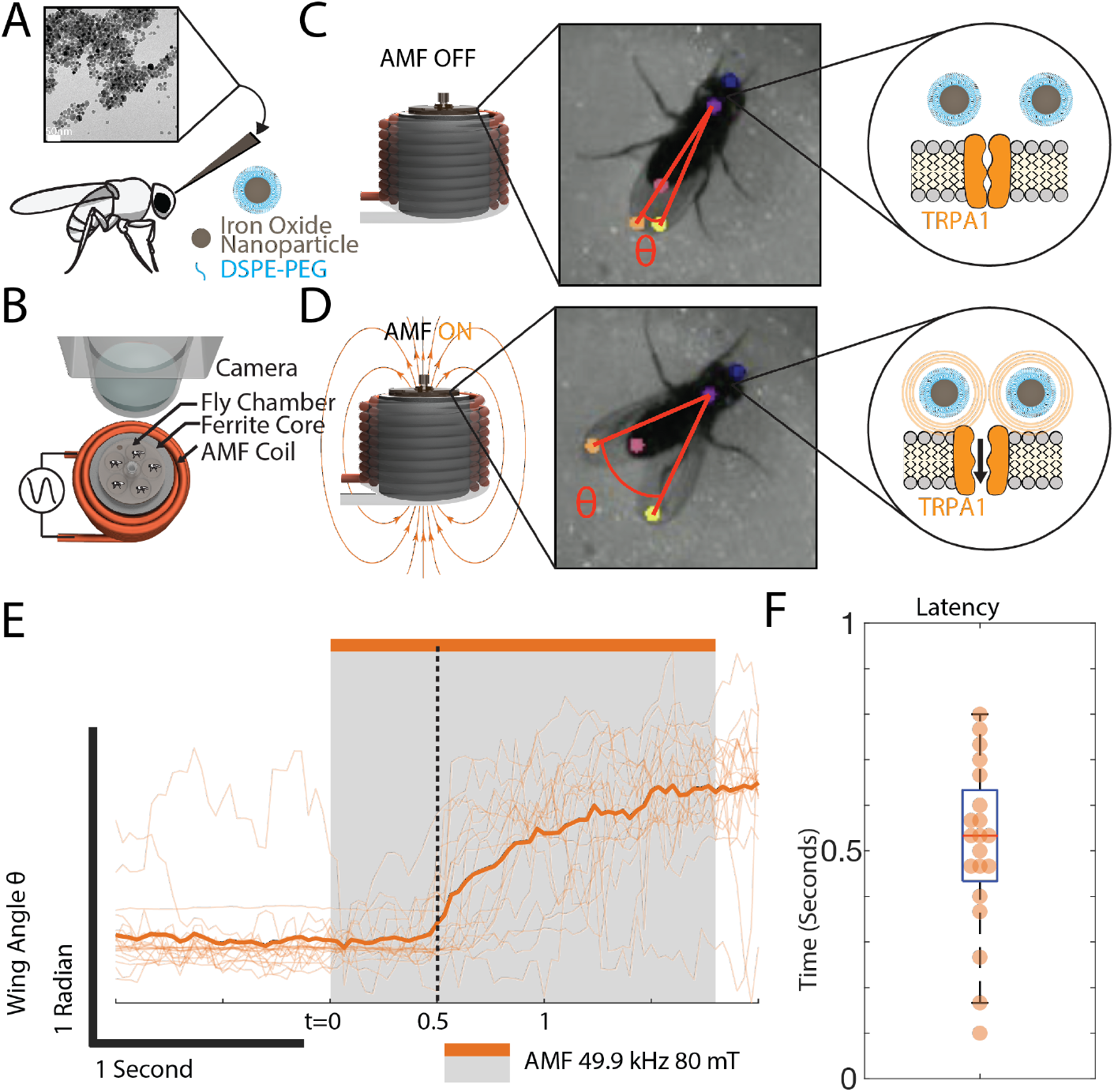
Behavioral Fly Assay. **A)** TEM picture and schematic of nanoparticle injection between the ocelli **B)** Freely moving flies in behavior chambers remotely stimulated by an induction coil and monitored by a camera to compare multiple animals at the same time. **C)** Placement of the behavior chamber on top of the magnetic coil with a ferrite core *(Left).* The flies are imaged at 30 fps to enable automatic posture estimation annotated by DeepLabCut *(Middle). Right* panel shows schematic nanoparticles in close proximity to TRPA1 channels. **D**) AMF activation of the coil *(Left)* results in a wing opening response *(Middle)* due to TRPA1-A channel activation by hysteretic heating of nearby nanoparticles *(Right).* **E)** Flies show distinct and reversible neuronal activation of cells expressing *fruitless* with sub-second behavioral responses repeatedly observed with the average of 5 flies showing responses to 4 repeated AMF stimulations (49.9 kHz; 80 mT) with **F)** an average behavioral response of 510 ± 186 ms as observationally determined by video *(**Video S2**).*

When we injected flies with 10 μg/mL of 15 nm cobalt-doped iron oxide nanoparticles and applied an AMF we observed a rapid increase in the wing angle with a response latency of 510 ± 186 ms observed from repeated stimulations on 5 flies — more than 10 times faster than previous *in vivo* magnetogenetic latencies^2,3^ (**Fig. 1E**). Conversely, the wing angle significantly decreases after ~370 ms and returns to baseline ~5.0 s after stimulation ends as calculated from the average of traces collected from 20 flies (**Fig. S9**). To confirm that the response was driven by magnetic heating of the nanoparticles, we compared wing openings in flies injected with 19 nm iron oxide particles to flies injected with 19 nm wüstite particles (**Fig. S4**). These wüstite particles contain the same amount of iron as the iron oxide particles, but with a smaller hysteresis loop, and thus do not heat well in an alternating magnetic field^28,29^ as characterized by AC magnetometry (**Fig. S5**). When we applied an alternating magnetic field to these flies, we observed a subsecond wing opening response in the iron oxide SPION-injected flies, but no response in the wüstite injected flies confirming that the behavioral response is mediated by magnetothermal heating (**Fig. S2**).

By comparing the fly response to fast and slow heating rates we were able to confirm that the wing opening response is indeed regulated by the rate-sensitive properties of TRPA1^17^. To perform behavioral experiments with different heating rates, we first characterized nanoparticle heating at two AMF conditions (49.9 kHz; 80 mT and 19 mT). These different AMF conditions result in a ~10× difference in heating rate (**Fig. 2A, B**), which are calculated as averages over the AMF duration due to visible lag from thermal resistance in the fiber optic probe. To assess the rate sensitivity of the behaving adult flies, we exposed the same set of flies to different temperature ramps by altering the magnetic field strength for different durations of time (1.8 s and 20 s; ΔT = ~O.82 °C *see methods)* while monitoring wing angle responses of adult males expressing TRPA1-A (**Fig. 2C, D**). The two heating conditions cause the tissue to reach a similar maximum threshold temperature but at different rates (**Fig 2. A, B**). The higher ΔT/t value of rapid heating lowers the threshold temperature of the TRPA1 channels^17^ and results in a statistically significant change in the wing opening phenotype (**Fig. 2F, G, H**). Assuming a similar thermal capacitance (C) for each fly, we expect all animals to receive the same total heat, but at different rates (ΔT = ΔQ/C). Experiments conducted on standard controls and flies expressing the temperature-threshold sensitive human TRPV1 channel show no response. This further demonstrates the improved sensitivity achieved by the rate sensitive TRPA1 channel (**Fig. S6, S7**).

**Figure 2.**
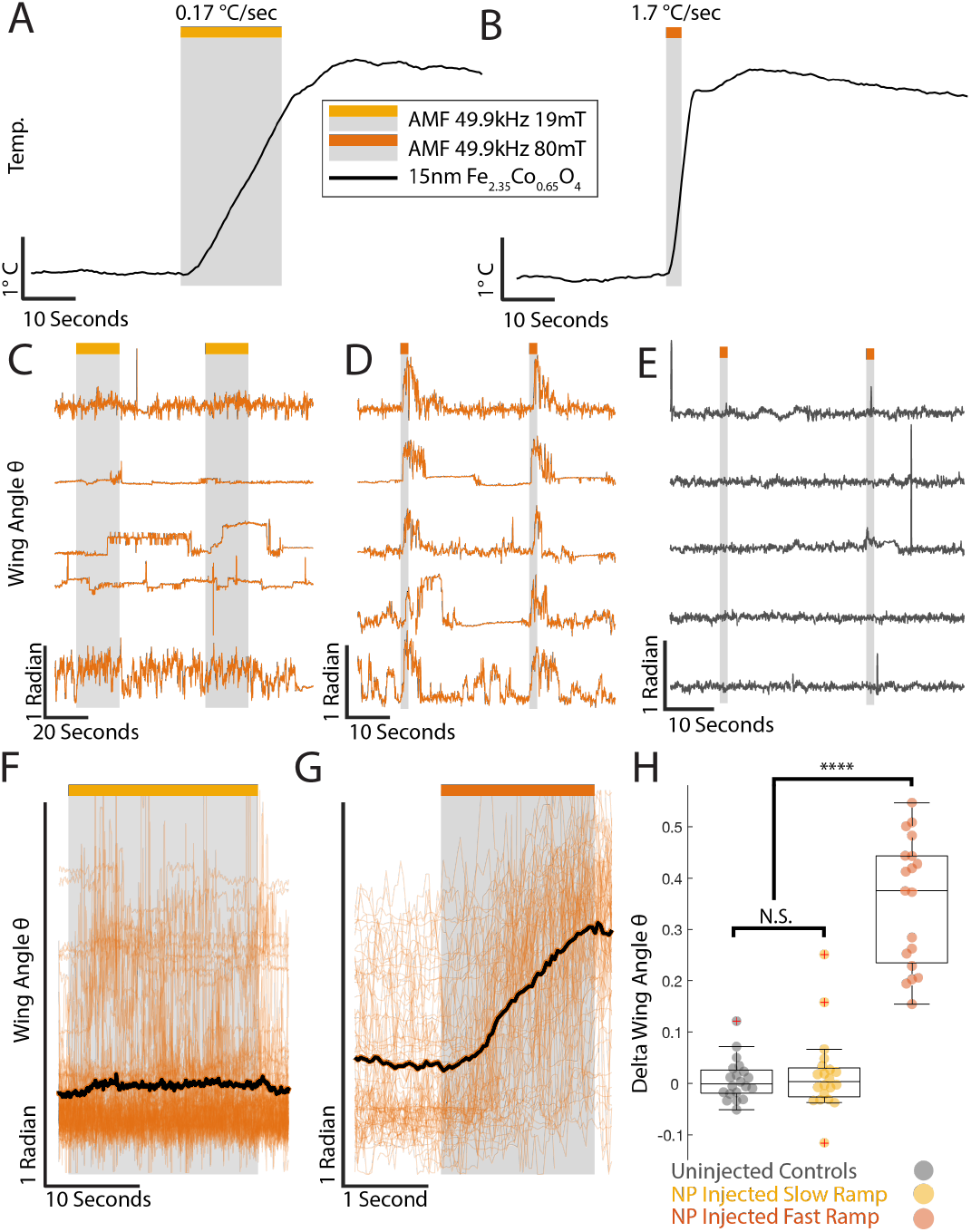
Rate response of magnetogenetic stimulation of cells expressing *fruitless* with sub-second response time. Thermal response of 15 nm cobalt-doped iron oxide nanoparticles at 10 mg/mL when exposed to alternating magnetic field for **A)** 20 s (49.9 kHz; 19 mT shown in yellow) or **B)** 1.8 s (49.9 kHz; 80 mT shown in red) reaching identical threshold temperature with rates of 0.17°C/s and 1.7 ^o^C/s respectively. Visible lag in heat measurement is due to thermal resistance in the fiber optic probe. **C-D)** Wing angle plots of the same 5 flies injected with 200 nL of 15 nm Iron Oxide nanoparticles (10 mg/mL) and expressing TRPA1 under the Fru-Gal4 driver when exposed to alternating magnetic fields for **C)** 20 s (49.9 kHz; 19 mT) or **D)** 1.8 s (49.9 kHz; 80 mT) and **E)** wing angle plots of uninjected flies exposed to 1.8s (49.9 kHz; 80mT). Stimulation experiments were repeated 2x for each stimulation protocol for the each set of 5 flies in the chamber **(Video S1, S2).** Average traces of 20 flies, each responding to 4 repeated AMF stimulations with **F)** Slow ramp (20 s; 49.9 kHz; 19 mT) and **G)** Fast ramp (1.8 s; 49.9 kHz; 80 mT). **H)** One-way ANOVA of delta wing angle taken immediately before and after stimulation and compared to uninjected control flies (**Fig. S3**) (**** = p< 0.0001). Post-hoc analysis with Tukey HSD Test. Normal distribution for each group determined by Chi-square goodness-of-fit test. For all data, outliers marked with red + if greater than [q3 + 1.5*(q3-q1)] or smaller than [q1 – 1.5*(q3-q1)] but still included in statistical analysis.

To generalize our findings, we also expressed the TRPA1 channel in a different neural circuit, producing a different behavioral effect. Driving expression of *dTRPA1-A* with *Hb9-GAL4* reliably induced a side-walking phenotype^23^ in the same behavior chamber under AMF control (**Fig. 3**).

**Figure 3.**
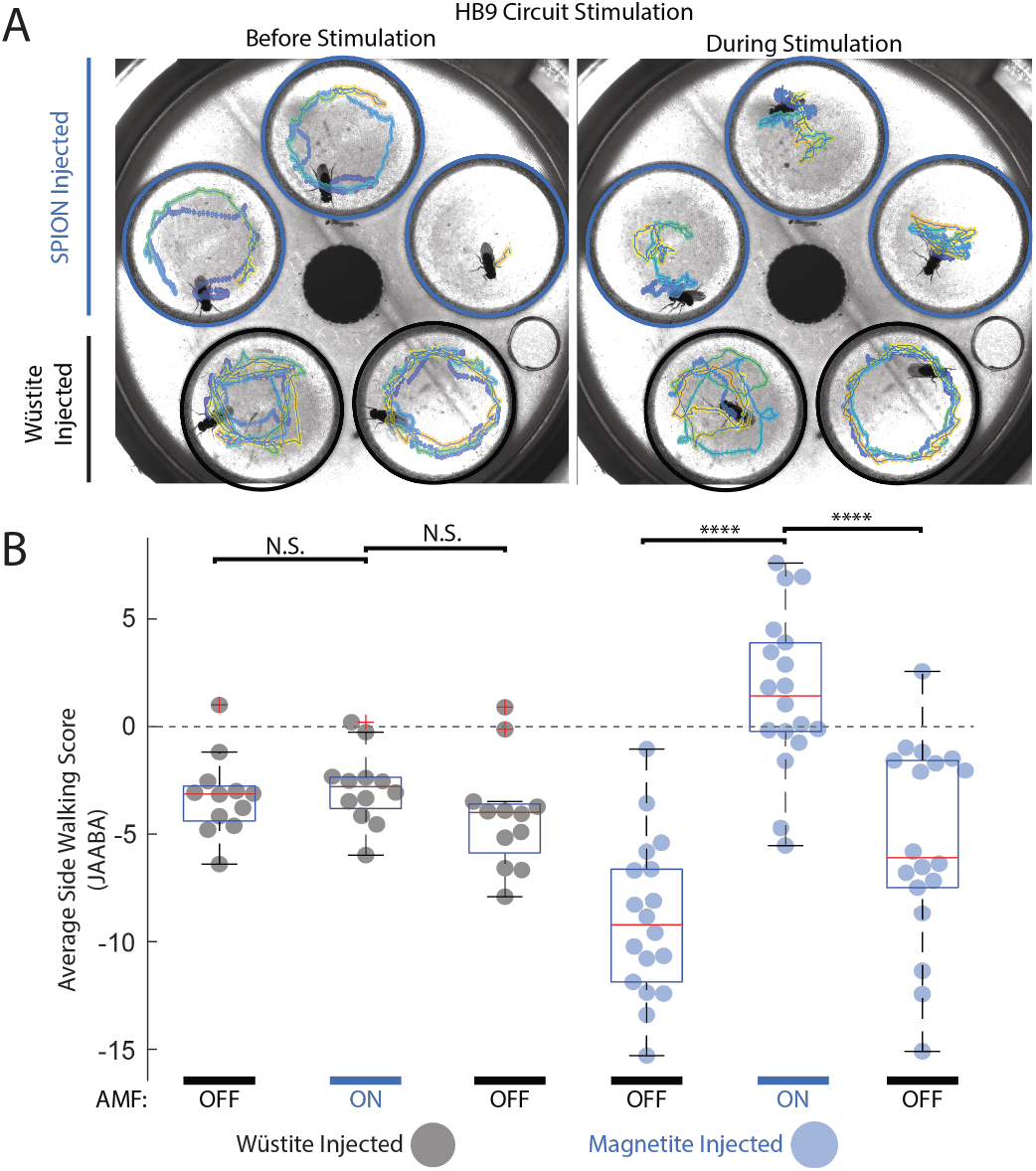
Versatility of magnetothermal stimulation in secondary cell type expressing Hb-9. **A)** Blue: Iron oxide SPION-injected, Black: Wüstite-injected controls. Purple to yellow gradient traces show 20 seconds of fly trajectories immediately before AMF stimulation and 10 seconds after the start of the magnetic stimulation (40 kA/m at 380 kHz). **B)** Box plot of average side-walking score from JAABA analysis of Magnetic stimulation of flies expressing TRPA1 under the control of the Hb9 driver trained by exogenous thermal stimulation of flies with the same genotype. Positive scores indicate likely side-walking behavior. Averages are taken over 20 seconds each immediately before stimulation, 10 seconds after the start of stimulation, and 10 seconds after the end of stimulation. N = 15 flies (9 injected with SPIONs; 6 injected with wüstite. One-way ANOVA and Tukey HSD test (**** = p < 0.0001) (**Video S3**).

We observed a clear, robust, and reversible side-walking phenotype during AMF stimulation among SPION injected flies expressing *dTRPA1-A* under the control of the Hb-9 driver. In contrast, flies injecting with poorly heating wüstite nanoparticles showed no response, demonstrating that the behavioral responses are due to specific nanoparticle heating and not an artifact of the magnetic field generation. The side walking behavior was more difficult to quantify than wing extension and took longer to develop. This increased latency may be due to the need to activate the peripheral nervous system where we expect fewer nanoparticles. Nevertheless, the magnetogenetic driven behavior was easily identified from the side-to-side behavior tracks (**Fig. 3A**) and the animal behavior videos (**Video S3**). FlyTracker generated a set of metrics for each track such as wing angle, velocity, angular velocity, and distance from the chamber wall. The combination of these FlyTracker metrics was used to train a machine learning model for side-walking behavior from videos for which side-walking behavior was hand annotated during thermal activation (35 °C) of flies expressing *dTRPA1-A* under the control of the Hb9-Gal4 driver using Janelia Automated Animal Behavior Annotator (JAABA)^30^. This model was then used to predict and annotate magnetically stimulated flies for similar behavior, providing a prediction score. Using FlyTracker and JAABA, we developed a classifier (see methods) to quantify the side-walking behavior before, during, and after stimulation. Quantification shows distinct and reversible modulation of side-walking behavior with iron oxide injected flies but not wüstite injected control flies (**Fig. 3B**).

We next explored whether magnetogenetic stimulation based on rate-sensitive thermoreceptors is compatible with reliable multichannel stimulation. By using nanoparticles that heat at different rates depending on the magnetic field conditions, we hypothesized that we could selectively activate flies injected with one type of nanoparticle (channel 1 – Ch1) without stimulating flies injected with another type of nanoparticle (channel 2 – Ch2), and *vice versa.* This is analogous to optogenetic stimulation of different neural circuits using different wavelengths of light, but here the selectivity is determined by differences in the specific absorption rate (SAR) of nanoparticles that we design and synthesize. This multiplexing concept is supported by the recent finding that modulating amplitude and frequency of an alternating magnetic field can selectively heat nanoparticles with varying coercivity resulting in multiplexed magnetothermal heating *in vitro^31,32^.* One limitation of the magnetic multiplexing modality demonstrated here is that although we can address nanoparticles independently, they activate the same ion channel (TRPA1). This technology is therefore best suited for targeting spatially segregated cell populations, either in different parts of the body or in different animals. The advantage of this type of multiplexing is that we can deliver a magnetic field that penetrates throughout a large volume of tissue (or multiple animals), and yet we can activate different spatially separated neuronal populations by changing the magnetic field strength and frequency.

To create the first channel for magnetogenetic heating we developed a highly coercive nanoparticle by doping iron oxide with cobalt (Fe2.35Co0.65O4) (**Fig. 4A**) which can generate a large amount of heat at a low frequency AMF with a high field strength (Ch1: 80 mT; 49.9 kHz). To create the second channel, we used a recently developed iron oxide nanoclusters^33^ (**Fig. 4B**) with a low coercivity to generate a large amount of heat when exposed to a high frequency AMF with a low field strength (Ch2: 12 mT; 555 kHz). These nanoparticles demonstrate a selectivity of ~15× for cobalt-doped iron oxide nanoparticles in Ch1 (SAR = 829.37 W/g for Cobalt-doped iron oxide; 50.57 W/g for Fe_3_O_4_ clusters) and ~10× for iron oxide nanoclusters in Ch2 (SAR = 31.60 W/g for Cobalt; 302.30 W/g for Fe_3_O_4_ clusters) when comparing heat generated over 3 s AMF stimulation (**Fig. 4D, E**) indicating 2 distinct channels available for magnetothermal heating. AC magnetometry further showed how coercivity and saturation differ between particles at relevant temperatures and AMF conditions (**Fig. S5**). Estimated nanoparticle SAR values from hysteresis loops measured in the double-sided coil varied slightly from measured nanoparticle SARs in single coil loop due to non-uniformity of magnetic fields. Crystal patterns for each particle were confirmed by X-ray power diffraction (XRD) with additional characterization of wüstite and iron oxide nanoparticles (**Fig. S4**).

**Figure 4.**
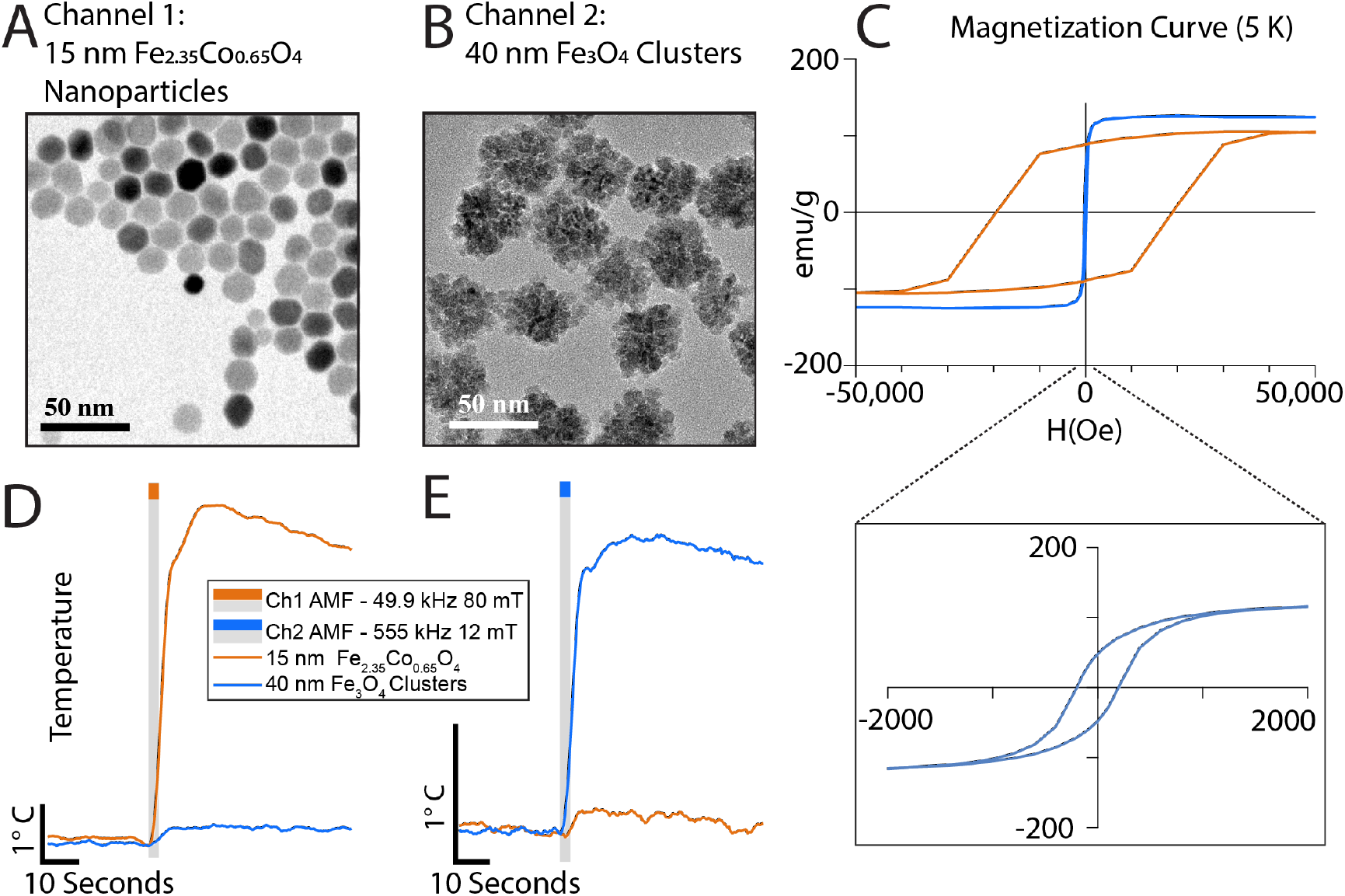
Multiplexed Magnetothermal Heating of Nanoparticles. **A)** TEM Characterization of 15 nm Cobalt-doped iron oxide nanoparticles. **B)** TEM Characterization of 40 nm Iron oxide nanoparticles clusters. **C)** Superconducting quantum interference device magnetometer data showing high variation in anisotropy between 15 nm cobalt doped iron oxide (Orange) and 40 nm iron oxide clusters (Blue) with inset showing more resolved hysteresis loop for low anisotropy clusters. **D-E)** Thermal response of 15 nm cobalt doped iron oxide nanoparticles (Red; 9.58 mg_metal_/mL) and 40 nm Iron Oxide nanoparticle clusters (Blue; 10.09 mg_metal_/mL) when exposed to an AMF of **D)** 49.9 kHz; 80 mT or **E)** 555 kHz; 12 mT.

When we measured wing angle in groups of flies injected with these different nanoparticles, we found that we could selectively drive wing extension in either group depending on which magnetic field stimulation channel we selected. Heating profiles of the nanoparticles were used to scale the injection concentration of each particle into fruit flies to achieve similar heating profiles in each magnetic field condition. We introduced a mixed group of flies injected with cobalt-doped iron oxide (orange circles) or iron oxide nanoclusters (blue circles) into the fly chamber and showed activation of behavior specifically in the optimized AMF and lack of a behavioral response in the off-target AMF condition (**Fig. 5A-C**). Exposure to 2 stimulations of Ch1 (2 s; 80 mT; 49.9 kHz) followed by 2 stimulations of Ch2 (2 s; 12 mT; 555 kHz) show the selectivity on animal behavior response based on injected particle while maintaining sub-second latencies for each channel (**Fig. 5B**).

**Figure 5.**
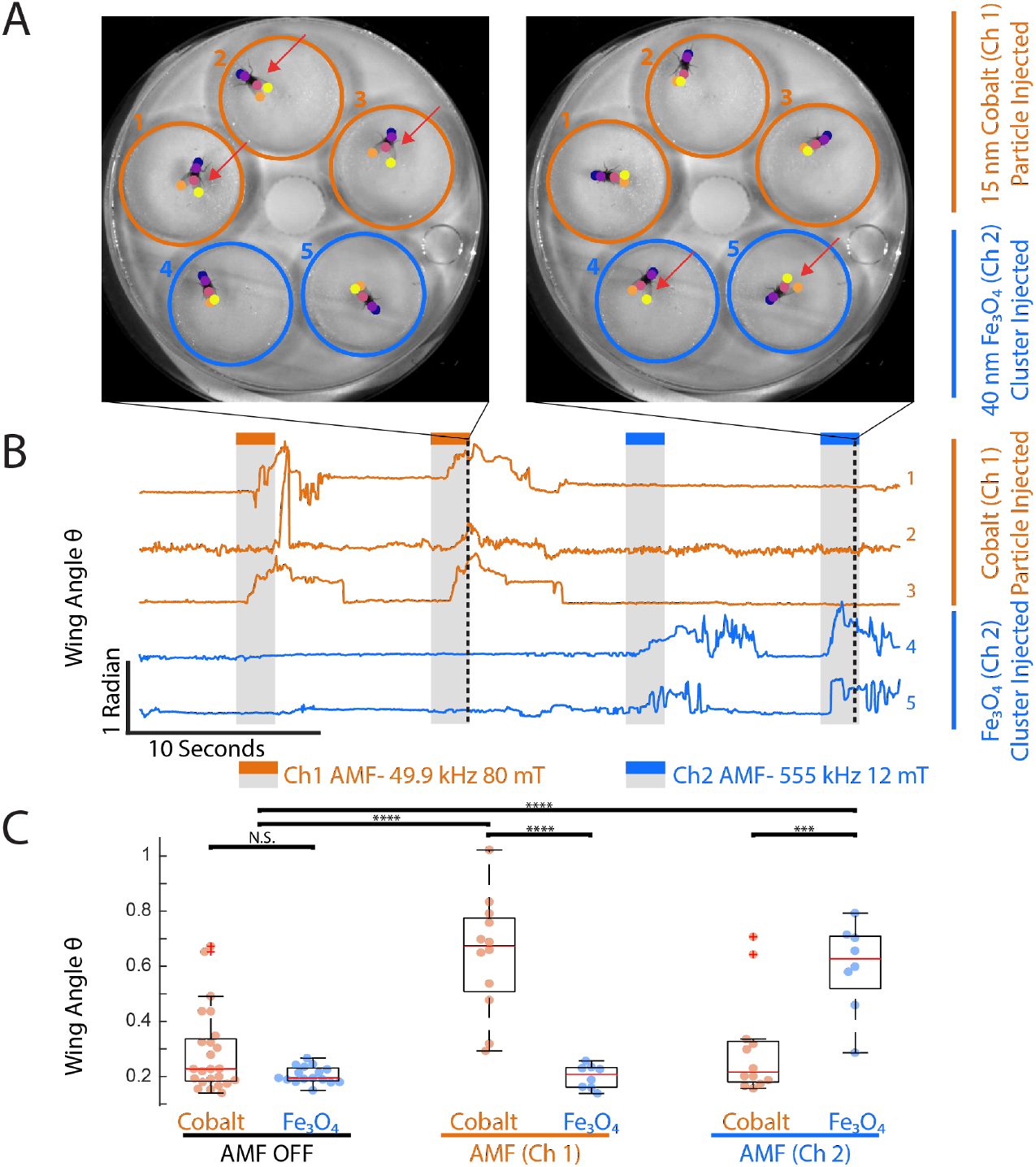
Multi-channel magnetogenetic stimulation of cells expressing *fruitless*. **A)** Example images of behavior chamber containing Drosophila with TRPA1-A expressed under the *Fru-Gal4* circuit selectively loaded with animals injected with 200 nL of 15 nm Cobalt Doped Iron Oxide (Top 3 chambers circled in red) or 40nm Iron Oxide Clusters (Bottom 2 chambers circled in blue). Left still is taken during an 80 mT 49.9 kHz AMF stimulus and shows wing extension of only the flies injected with 15 nm Cobalt doped particles. Right still is taken during 12 mT 555 kHz AMF stimulus and shows wing extension of only the flies injected with 40 nm iron oxide clusters. **B)** Wing angle plots tracked by DeepLabCut of 5 flies shown when exposed to 2 s pulses magnetic fields (Red: 49.9 kHz; 80 mT or Blue: 555 kHz; 12 mT). Top 3 fly traces are flies injected with 200 nL of 15 nm Cobalt Doped Iron Oxide while bottom 2 traces are flies injected with 40 nm Iron Oxide Clusters. **C)** Box plots with scatter plots showing the wing angle of each fly immediately before (AMF Off) and after 2s of magnetic stimulation (Ch1 and Ch2). N = 5 flies. One-way ANOVA (*** = p< 0.001; **** = p< 0.0001) **(Video S4)**

In summary, we report the first multiplexed magnetothermal activation of behavior in freely moving adult *Drosophila melanogaster,* and the first sub-second magnetogenetic response *in vivo.* This subsecond response was made possible by replacing the slow-response magnetothermal sensor TRPV1 by a rate-sensitive TRPA1 channel. *Drosophila* TRPA1-A is rate sensitive and native to flies. The magnetic activation of the channel drives behavior *in vivo* within 500 ms of stimulation for which we estimate thermal temperature increases in the tissue to be less than 1 °C based on nanoparticle heating and the average mass of adult male *Drosophila* (see *Methods*). Thermal imaging *(FLIR A700)* confirmed no significant heating of the surface of the fly (**Fig. S11**) as well as no notable heating of the chamber during magnetic stimulation (**Fig. S10**).

Future applications with targeted nanoparticles may enable multiplexing with similar channels with a heterogeneous population of target neurons or cells within the same volume. However, due to the size of the *Drosophila* nervous system, heat transfer limitations^34^, and the thermal rate required to activate these channels to direct behavior^17^, highly concentrated ferrofluids show the most promise for current neuronal stimulation applications. Further sensitization and optimization of the thermal rate response may make heterogeneous multichannel targeted activation of the nervous system possible by genetically targeting the nanoparticles to bind to specific membranes or channels.

The ideal magnetothermal sensor for mammalian stimulation at 37 °C may be found among the orthologous TRPA1 proteins in reptilian or avian species and/or through protein engineering, including site-specific mutagenesis or protein chimerization. This new magnetogenetic method depends on the *Drosophila* dTRPA1-A which is constitutively active at 37 °C. In order to adapt this approach to stimulate mammalian neurons, other channels with similar temperature rate sensitivities but higher threshold must be characterized or engineered. The heat responses of reptilian and avian TRPA1 channels have been described showing a conserved heat response in animals like the western clawed frog, chicken, green anole, rat snake, and rattlesnake^35–37^, which might be promising candidates. These thermal responses have further proven to be heavily reliant on the ankyrin repeat N-terminus domain in both *Drosophila* and snakes^17,38^, which therefore constitute an ideal target for future protein engineering.

With the relatively fast response and multiplexing abilities of magnetogenetics shown here we believe that this technology has the potential to rival optogenetics in terms of temporal resolution and multiplexed stimulation while maintaining the advantages of remote activation over large volumes of cells that may lie in deep tissue such as brain tissue occluded by the skull.

## Methods

### Generation of Biocompatible Magnetothermal Transducer and Control Particles

#### Superparamagnetic Iron Oxide (Fe_3_O_4_) Nanoparticle synthesis

As iron oxide nanoparticles has shown promising biocompatibility^39^, we synthesized super-paramagnetic iron oxide nanoparticles (SPIONs) consisting of iron oxide nanocrystals and coated with a layer formed of copolymers of phospholipids and poly(ethylene glycol) (DSPE-PEG2K). The nanoparticles were synthesized, coated and functionalized in three consecutive steps similar to previously published work^40^. First, 4 nm iron oxide nanocrystals were synthesized by thermal decomposition of iron acetylacetonate in a mixture of oleic acid and benzyl ether. The iron oxide nanocrystals were then grown to 19 nm diameter by controllable seed-mediated growth in a mixture of iron acetylacetonate, oleic acid, and benzyl ether. The size distribution of the iron oxide nanocrystals was then quantified by transmission electron microscopy (TEM). The magnetic properties and the crystal structure of the nanocrystals was then quantified by a superconducting quantum interference device (SQUID) and power X-ray diffraction (XRD).

The synthesized nanocrystals were then coated with a layer of oleic acid and only dispersible in a nonpolar solvent. To generate water-dispersible nanoparticles, the nanocrystals were coated with a mixture of DSPE-PEG2K using a dual-solvent exchange method^41^. The hydrodynamic size of conjugated nanoparticles was subsequently examined by dynamic light scattering. The heating efficiency of SPIONs was then examined by a magnetic inductive heating within the AMF device using a fiber optic thermal probe (Lumasense Luxtron 812 and STF-2M Probe).

#### Cobalt Doped Iron Oxide (Fe_2.35_Co_0.65_O_4_) Nanoparticle synthesis

The cobalt-doped iron oxide nanoparticles were made by multiple seed-mediated growth reactions using 5nm iron oxide cores. They were synthesized through thermal decomposition, using 2 mmol CoCl_2_, 4 mmol Fe(acac)_3_, 25 mmol oleic acid and 60 ml benzyl ether as a solvent. The reaction was heated to 120 °C for 30 minutes under a constant argon flow, then to 200°C for 2 hours and finally to reflux at 300 °C for 30 minutes. The product was purified through several acetone washes. The nanoparticle sizes were determined by HC TEM. Nanoparticles were then coated with DSPE-PEG2K by mixing the nanoparticles with PEG and adding DMSO. The reaction was then evaporated and transferred to water by a drop-wise addition of water and removal of the remaining DMSO by was done by centrifugation and ultracentrifugation.

#### Iron oxide (Fe_3_O_4_) nanocluster (40 nm) synthesis

The 40 nm iron oxide nanocrystal clusters were synthesized through the hydrothermal reaction. FeCl_3_·6H_2_O (540 mg) was dissolved in ethylene glycol (20 mL) under vigorous magnetic stirring. Then poly(acrylic acid) (250 mg), urea (1200 mg), and ultra-high purity deionized water (1.0 mL, < 18 mΩ) were added to the solution. The mixture was vigorously stirred for 30 min, yielding a transparent and bright yellow solution. The mixture was then transferred to a Teflon-lined stainless-steel autoclave, tightly sealed and then heated at 195 °C for 6 hours with a temperature ramp rate of 20 °C/min. After the reaction mixture was cooled to room temperature, the product was collected using a magnet. The clusters were washed 6 times using ethanol and water to remove the unreacted reactants and byproducts and then dispersed in DI water. The sizes of clusters and primary particles were determined using TEM. More than 500 clusters were measured to determine the cluster dimensions.

#### Wüstite (FeO) nanoparticle synthesis

Synthesis of the control wüstite (FeO) nanocrystals, which are poor magnetothermal transducers, was achieved similarly to the SPION synthesis with minor alterations. Benzyl ether was substituted with oleylamine, the initial reaction was lengthened with a reduced temperature, and a vacuum process was added. The wüstite nanocrystals were then purified with ethanol, surface treated via heating in oleylamine, and dispersed in toluene before they were coated with DSPE-PEG2K.

The samples for TEM measurements, both HC TEM and TITAN TEM, were prepared by diluting the samples and placing them in carbon-film grids. XRD samples were prepared by drying the nanoparticles under an argon flow and then pulverizing the resulting powder. SQUID measurements were done with coated samples by fixing the nanoparticles with calcium hemisulfate and enclosing them within a capsule to prevent movement and normalized in terms of metal as determined by ferrozine assay and ICP. Doping percentages were determined by ICP-MS and the samples compared to the corresponding standard curves of iron and cobalt. Specific absorption rate (SAR) is calculated as:

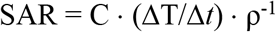

Where C is specific heat capacity of the media (C = 4180 J·kg·K^-1^), T is the total temperature change during stimulation averaged over 3 stimulations, *t* is the AMF stimulation time and ρ is sample density measured by total metal concentration. Iron oxide nanoclusters were recorded at 10.09 mg_metal_/mL and cobalt-doped iron oxide nanoparticles were recorded at 9.58 mg_metal_/mL. Temperature was measured with a fiber optic thermal probe (Lumasense Luxtron 812 and STF-2M Probe) which is unaffected by magnetic fields.

### Fly stocks and husbandry

Parental *Drosophila* strains were obtained as a gift from the Venkatachalam lab UAS-hTRPV1 P{w[+mC]=UAS-VR1E600K}(Strain generated by random P-element insertion and maps to the second chromosome)^42^

> or from the Bloomington Drosophila Stock Center:
>
> *Fru-GAL4* (BL66696) w[*]; TI{GAL4}fru[GAL4.P1.D]/TM3, Sb[1]^43^ *UAS-TrpA1-A* (BL26263) w[*]; P{y[+t7.7] w[+mC]=UAS-TrpA1(B).K}attP16^44^ *Hb9-GAL4* (BL32555) w[*]; P{w[+mW.hs]=GawB}exex[Gal4]

P{w[+mC]=lacW}nsl1[S009413]/TM3, P{w[+mC]=GAL4-Kr.C}DC2, P{w[+mC]=UAS-GFP.S65T}DC10, Sb[1]^45^.

All flies were reared on cornmeal, molasses, sugar, yeast, and agar food, on a 16 h light/8 h dark cycle, and at room temperature (22.5 ± 0.5 °C).

### Nanoinjection of Nanoparticles into *Drosophila* Heads

Different *GAL4* driver lines were crossed to *UAS-TRPA1* flies and offspring with both *GAL4* and *UAS* components and single component controls were collected for injection. Nanoparticles were injected into adult male heads similar to a previously described protocol^46^. Males that were 1 to 5 d old were immobilized on ice and dropped head-down with an aspirator into a cylindrical hole punched with a pasteur pipette tip into a 2% ice-cooled agarose gel approximately 5mm thick. The flies were then aspirated through the gel until the top of their head was flush with the gel surface. Five flies were immobilized in a gel at a time and transferred to a thermoelectric temperature controller (TE Technology Inc., Traverse City Michigan). Using a Nanoject II (Drummond Scientific Company), and borosilicate needle pulled on a Model P-97 needle puller (Sutter Instruments), nanoparticles resuspended in artificial *Drosophila* hemolymph^46^ were aspirated into the needle. Using a micromanipulator (Narishige, Model M152) attached to a fixed post to move the Nanoject in 3 dimensions, the needle tip was placed just above the top of the fly head sticking out of the refrigerated gel and positioned between the 3 ocelli at a 45° angle. The flies were then injected by gently pushing the needle forward until it penetrated the cuticle between the ocelli. Approximately 200 nL of nanoparticles suspended in artificial hemolymph were injected directly into the brain and flies were aspirated through the gel into an empty vial containing standard fly food. Iron oxide nanoparticles of 19 nm diameter were injected at 10 mg/mL, cobalt doped iron oxide particles were injected at 10 mg/mL, and 40 nm iron oxide clusters were injected at 25 mg/mL. Animals were then allowed to recover overnight before being placed in a behavior chamber and stimulated with AMF (**Figure 1**).

### AMF Stimulation of *Drosophila*

Animals were given at least 16 hours to recover from nanoparticle injection before being loaded into the AMF generator. Experimental and control animals were each placed into one of five cylindrical arenas (12 mm diameter) in the behavioral chambers within a 50 mm diameter enclosure by aspiration through a small hole cut into an acrylic cover that can rotate over each arena. This 3D printed behavior chamber is then placed into a 3D printed chamber holder which places the animals in the center of a 17 turn 50 mm inner diameter (ID) coil (Nanotherics Magnetherm) for the courtship behavior using 19 nm SPIONs (Figure S2) or a custom high powered 6 turn 50 mm ID coil (Fluxtrol/AMF Lifesystems) for sidewalking behavior experiments via inductive heating. Stimulation of cobalt nanoparticles and iron oxide nanoclusters were performed by placing the chamber ~6 mm above the surface of a 6-turn 57.7 mm ID Hi-Flux coil with a ferrite core (μ = 2300) (MSI Automation) driven by a custom FPGA-controlled hybrid Silicon–Gallium-Nitride transistor based power electronics system (Duke University), which can generate AMF in the same coil at several distinct frequency channels spanning 50 kHz to ~5 MHz and rapidly switching between the channels on millisecond time scale. The camera (Basler acA2000-165umNIR, 50mm F1.8 Edmund optics 86574) was then fixed above the animals and synced with the alternating magnetic field via TTL triggers to temporally align behavioral recordings with magnetic field generation. Frequency was set by the machine while field strength was measured by a magnetic field probe placed in the same location as the fly behavior chamber (Fluxtrol).

Thermal ramp demonstration used 2 stimulations of 1.8 s duration at 80 mT and 49.9 kHz for fast heating and 20 s duration at 19 mT and 49.9 kHz for slow heating. The inter stimulation intervals were 30 or 60 *s* for fast and slow heating respectively. Multichannel demonstration used exposure to 2 stimulations of Ch 1 (2 s; 80 mT; 49.9 kHz) followed by 2 stimulations of Ch2 (2 s; 12 mT; 555 kHz) with inter stimulation interval of 10 s. The video recording is paused for less than 1 second in the multiplexing recording (t=20 *s*) to switch the stimulation protocol on the software from Ch1 to Ch2.

### Automated Analysis of Behavioral Phenotypes in *Drosophila*

Animals were given at least 5 minutes to adjust to the behavior chamber before stimulation with pulsed cycles of alternating magnetic fields. Backlit videos of flies were analyzed using the Caltech FlyTracker^27^ to automatically identify the fly position, orientation, and wing/leg extensions. These data were then be analyzed on a frame-to-frame basis in MATLAB (MathWorks) for specific phenotypes (e.g. maximum wing angle for the wing extension phenotype and fly position for side-walking phenotype) or used with machine learning tools like Janelia Automated Animal Behavior Annotator (JAABA) to train complex behaviors that take place over a series of frames (lateral movement phenotype in side-walking) which enables linear regression models to predict the occurrence of the phenotype. Multiplexed animal behavior was analyzed using DeepLabCut^26^ for dynamic tracking of flies with visible shadows introduced by front lighting needed to illuminate flies above the ferrite core. The animal analysis was trained using a skeleton labeling the head, neck, tip of each wing, and abdomen. Wing angle was calculated between the neck and each wingtip.

Each experiment consisted of 2 AMF stimulations per fly and was repeated twice per fly. Experiments were performed on a minimum of 20 flies for control groups and 40 flies for those expressing TRPA1 under the fruitless driver. Traces were sorted by average area under curve during magnetic stimulation and the top 10 flies from each group were used to calculate comparisons from control vs experimental groups (**Fig, S6, S7**) while the top 20 flies were used to compare across conditions for flies expressing TRPA1 under the fruitless driver (**Fig. 2**). The selection of the most responsive 10 or 20 flies from each group was done to remove data from flies that were poorly injected, which is a known challenge related to backflow of the nanoparticle solution. Additionally, this selection process eliminates data with DLC tracking errors. Power analysis performed with G*power on an average of 5 flies suggested sample sizes of n=5 is sufficient for experimental against control groups and n=8 is sufficient for thermal rate comparisons against controls (**Fig. S8**).

Side walking and multiplexing experiments were done with fewer flies as these experiments show extensions of the basic experimental approach. Statistics for **Fig. 3B** are shown for each stimulation with N=15 flies (9 injected with SPIONs and 6 injected with wüstite with 2 repeated stimulations for each fly. Statistics for **Fig. 5C** are shown as individual simulations of each fly N = 6 cobalt injected, N = 4 nanocluster injected with 4 repeated stimulations for each channel for each fly.

### Thermal Imaging of Fly Chamber and Immobilized Flies

To assess thermal stability of the chamber during magnetic stimulation, the fly chamber put into position above the magnet without the acrylic lid to enable thermal imaging of the interior of the fly chamber. Previously injected flies expressing TRPA1 under the control of the *fruitless* gene that were shown to be responsive were immobilized and placed into the chamber. Thermal imaging (FLIR A700) was performed on the chamber and flies to verify ambient heat. Regions of interest were traced around the fly and analyzed using manufacturer’s software (FLIR research studio). Raw traces of individual chambers without flies under magnetic stimulation is shown (**Fig. S10**) and fly heating data is shown by subtracting the temperature of a nearby area within the chamber without an injected fly to offset baseline fluctuations introduced from the camera sensor (**Fig. S11**).

### Theoretical Calculation of Thermal Fluctuation of Injected *Drosophila*

To determine the thermal fluctuation of the tissue in the fly during magnetic stimulation, we can use the density-based SAR of the nanoparticles to get the delta temperature of the mass of the *Drosophila* using a simple mass dilution. This calculation can be used to get a rough estimate of the expected temperature fluctuation of the thermal mass within the animal. For this measurement we assume the mass of the adult male fruit fly is ~0.88 mg as previously shown^47^ and has a specific heat capcity similar to water (4180 J·kg^-1^·K^-1^).

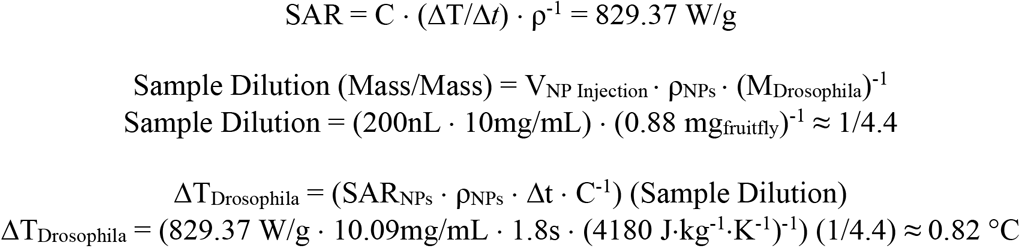

As this calculation relies on assumptions of thermal capacitance (C) of each fly being approximately the same as water, we expect that animals in all experiments receive the same total heat but at different rates (ΔT = C/Q). As such, we simply estimate an upper bound of ~1 °C overall change in the thermal temperature of the bulk fly tissue. While we suspect that there could be localized areas within the fly that experience more or less temperature change with nonuniform particle distribution, tissue damage from hyperthermia can be reasonably ruled out.

### AC Measurement of Nanoparticle Dynamic Magnetization

A custom double-sided, high-amplitude alternating magnetic field (AMF) generator was constructed from super-conductive copper tubing (10 AWG equivalent) and two E-shaped, N87 ferrite cores (μ = 2200; TDK Electronics). The cores were each wrapped by 9 turns, then assembled with material between the outside arms to create an air gap of 5.3 mm between the middle arms. The two coils and a resonant capacitor were wired in series, and the circuit was driven by a custom air-cooled Gallium-Nitride transistor-based power electronics board (Duke University). This driver board consisted of an H bridge powered by a voltage-controlled DC power supply (Aim-TTI QPX1200S) and gated by a two-channel function generator (BK Precision 4052).

A 17 μL sample of nanoparticles (~10 mg/mL) suspended in water was loaded into a 3D-printed, hollow chamber. The chamber was sealed with Scotch tape and placed onto a custom two-layer, 16 mm-thick alternating current magnetometer (ACM) circuit board (MIT)^31^. The ACM board was then positioned within the AMF generator air gap such that the field was completely uniform across the board’s two oppositely-wound, inductive pickup coils (one containing the sample and the other vacant). Field strength measurements from the circuit board’s single-turn pickup coil were calibrated with an HF magnetic field probe (Fluxtrol). At each field strength, the AMF generator was driven for 200 ms at 55 kHz and the last 100 periods of the signals induced by the applied AMF and the changing nanoparticle magnetization were captured, filtered, and amplified on the ACM board. The same protocol was run with 17 μL of water in the sample chamber.

The magnetization signals from the nanoparticle and water samples were subtracted to further reduce noise and isolate the true nanoparticle magnetization. The resulting voltage signals were integrated to yield the applied magnetic field and nanoparticle dynamic magnetization signals. The magnetization was then normalized with respect to sample concentration, then calibrated by setting the saturation magnetization to that measured by SQUID. The 100 collected periods were each centered and the average was taken across all periods to produce the average hysteresis loop.

## Supporting information

Supplemental Video S1 | Slow Thermal Ramp of Cobalt Injected Drosophila for Fru circuit stimulation

Supplemental Video S2 | Fast Thermal Ramp of Cobalt Injected Drosophila for Fru circuit stimulation

Supplemental Video S4 | Multiplexed Drosophila Fru circuit stimulation

## Acknowledgments

This research was developed with funding from the Defense Advanced Research Projects Agency (DARPA) of the United States of America, Contract No. N66001-19-C-4020. The views, opinions and/or findings expressed are those of the authors and should not be interpreted as representing the official views or policies of the Department of Defense or the U.S.A. Government.

This work was funded in part by the National Science Foundation – NSF Neuronex innovation award 1707562, grant C-1963 from the Welch Foundation, and National Institutes of Health under award number RO1MH107474.

We also thank Junsang Moon and Polina Anikeeva (MIT) for useful discussions and guidance with magnetic multiplexing.

## Notes

### Competing Interest Statement

S. M. Goetz has received research funding from Magstim. A. V. Peterchev has received research funding, travel support, patent royalties, consulting fees, equipment loans, hardware donations, and/or patent application support from Rogue Research, Tal Medical/Neurex, Magstim, MagVenture, Neuronetics, BTL Industries, and Advise Connect Inspire.

### Summary of Updates

Updated supplemental information (Supplemental Figures 5-11) with TRPV1 comparison, more controls, and more statistics for groups. Thermal calculations and AC Magnetometry data has also been added.

