## Supplemental Figures and Video Legends for "Sub-second multi-channel magnetic control of select neural circuits in behaving flies"

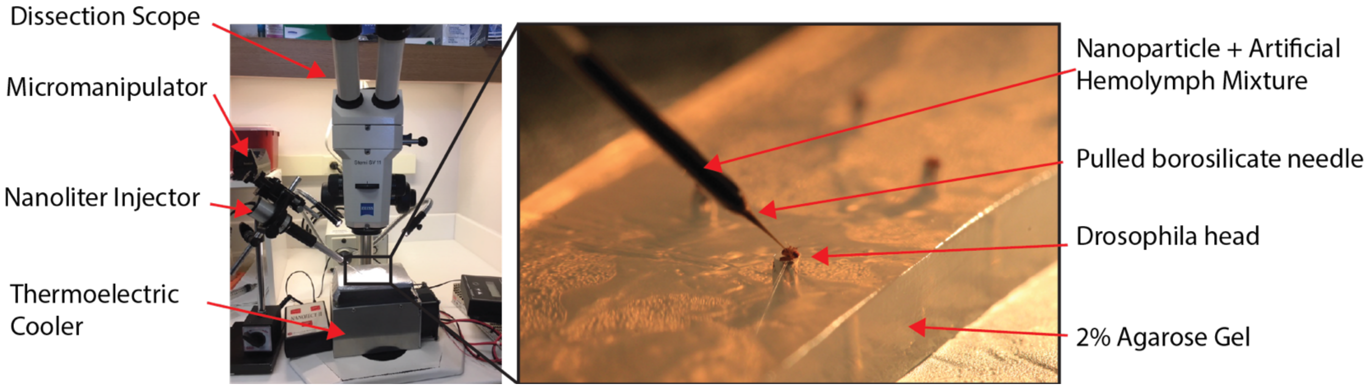


**Supplemental Figure S1 | Drosophila nanoinjection set up.**
Left: Macro view of nanoinjection apparatus below a dissection scope. Right: Cold immobilized *Drosophila* loaded into a 2% Agarose gel for injection between the ocelli.

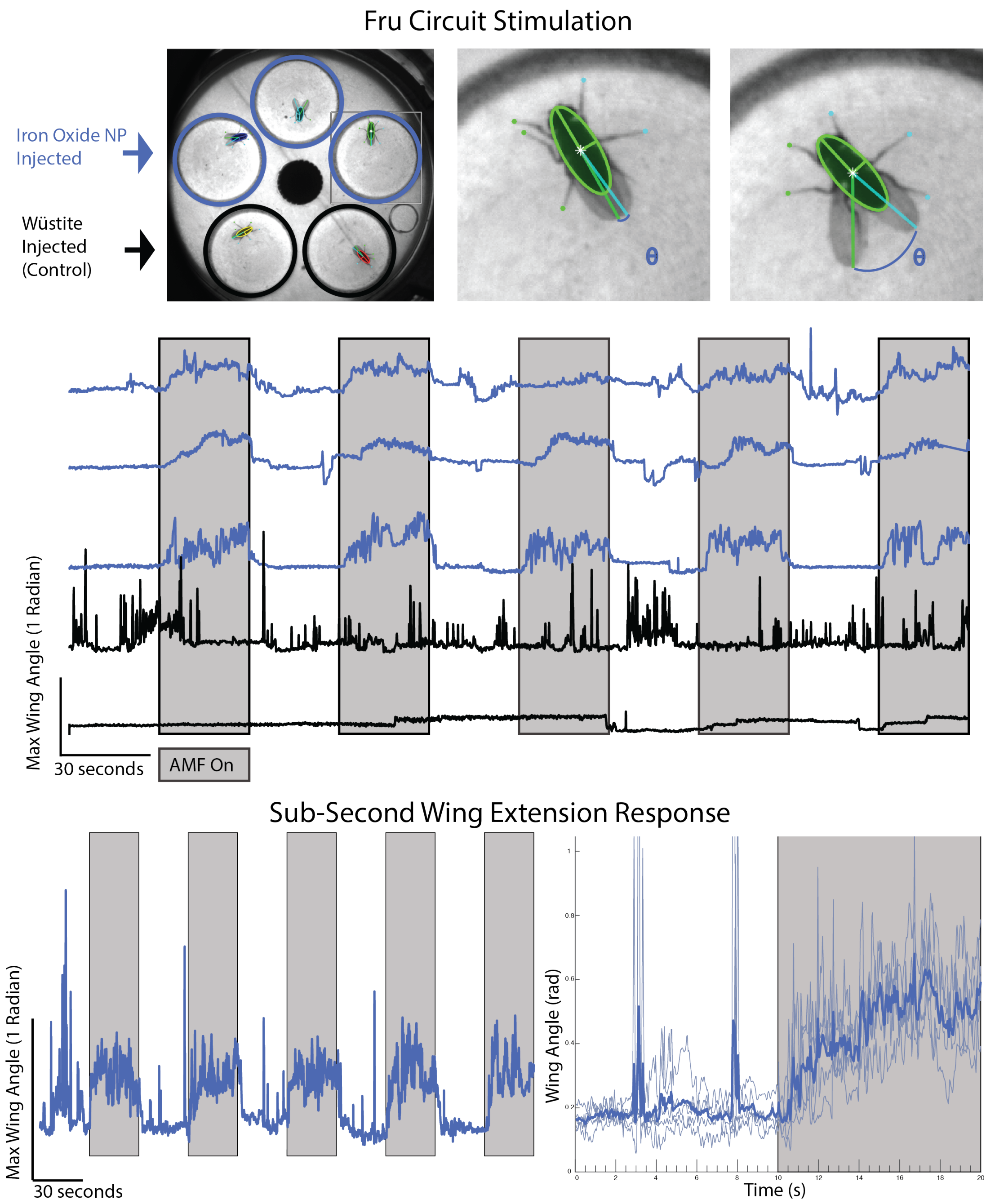


**Supplemental Figure S2 | TRPA1 magnetogenetics drives *fast* behavioral responses in freely behaving flies.** Top: *FlyTracker* analysis of fruit fly wing extension. Middle: Traces of wing angle (FlyTracker) when exposed to 30 s pulses of AMF stimulation (10 kA/m; 390 kHz) shows robust response in 19nm Iron Oxide ($Fe_3O_4$) particle injected flies (blue) with a specific absorption rate 245 W/g when compared to non-heating 19nm wüstite (FeO) injected controls (black). Bottom: Example fly data shows sub-second behavioral wing extension in response to AMF stimulation. All experimental repeats (N=5; thin lines) are averaged (bold line).

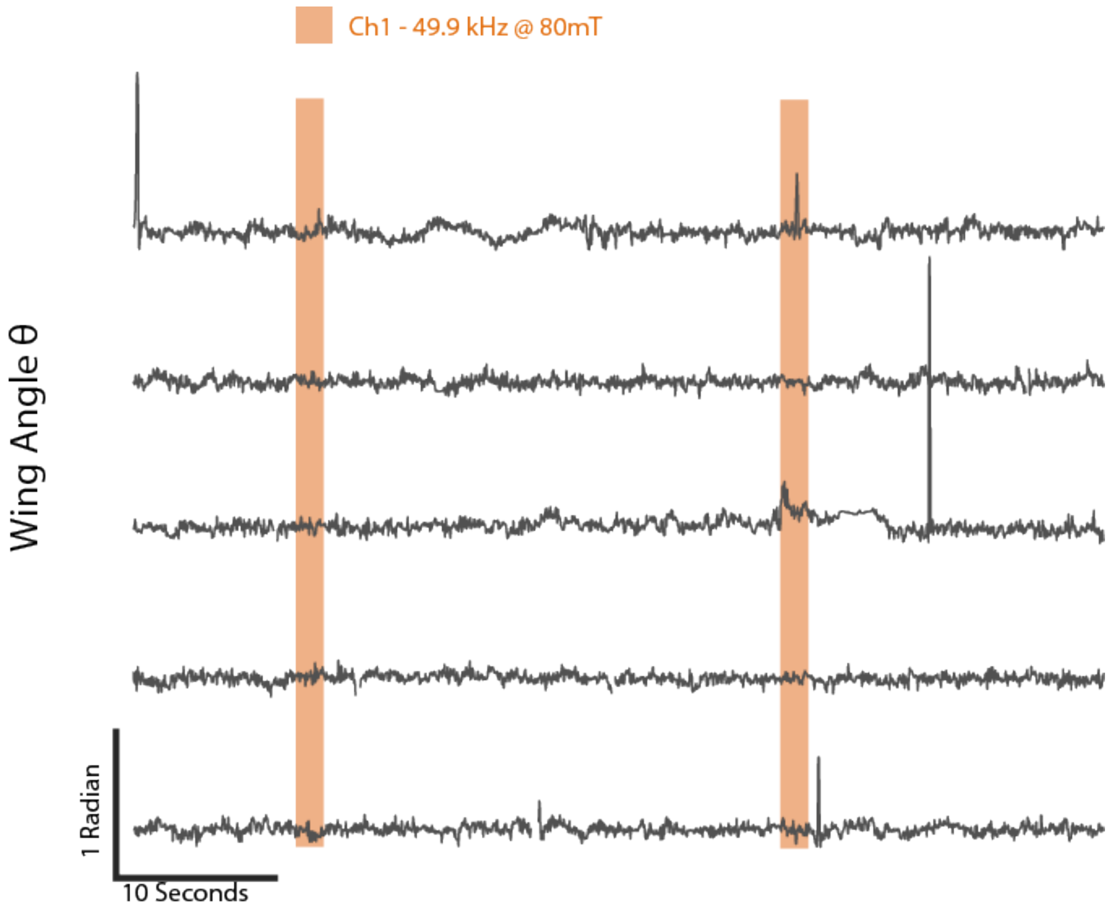


**Supplemental Figure S3 | Uninjected Control Fly Response to AMF stimulation.**
Example wing angle measurements quantified using DeepLabCut for 5 uninjected flies expressing *TRPA1* in *Fru-Gal4* circuit when exposed to alternating magnetic fields for 2 stimulations of 1.8 s (49.9 kHz; 80 mT). No significant wing opening response during AMF is observed.

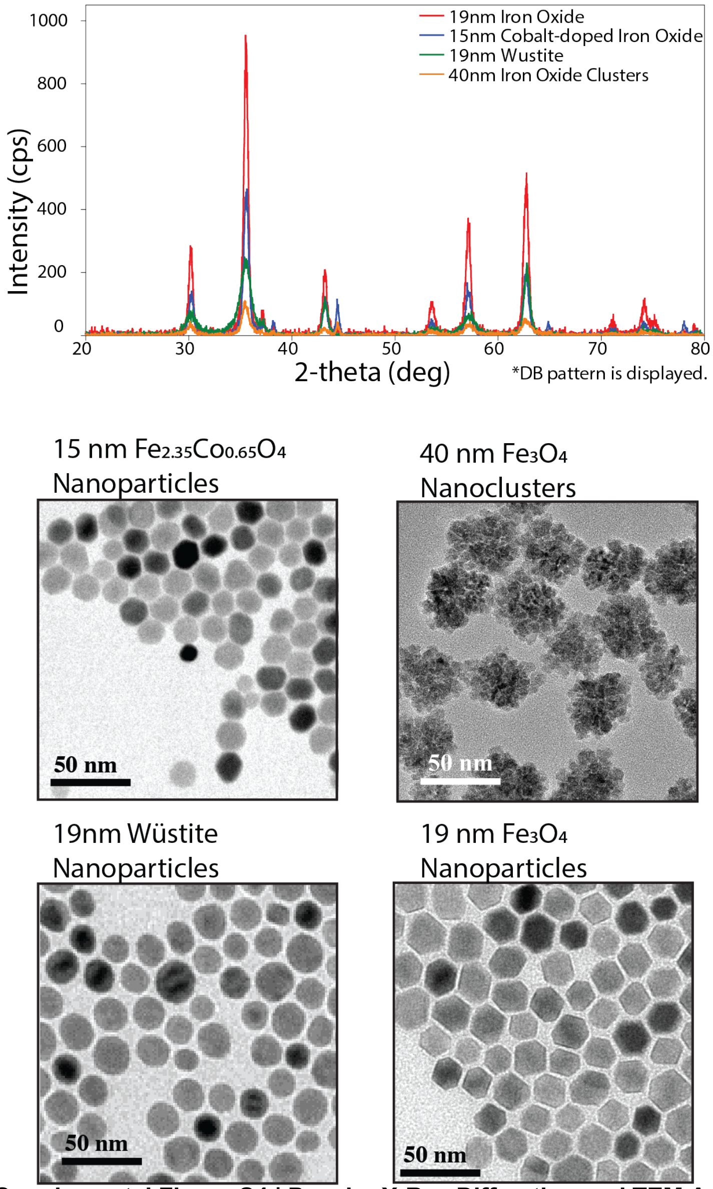


**Supplemental Figure S4 | Powder X-Ray Diffraction and TEM Analysis of Nanoparticles.**
X-Ray diffraction analysis using Cu Ka radiation, vertical theta/theta goniometer, graphite monochrometer and scintillation counter. Overlayed pattern of the samples: $Fe_3O_4$ nanoparticles (red); $Co_{2.35}Fe_{0.65}O_4$ nanoparticles (blue); Wustite (FeO) nanoparticles (green); $Fe_3O_4$ nanoparticle nanoclusters (oeange). Magnetite NP showed a crystallite size of 14.9 nm and a lattice parameter of 8.3836 Å, cobalt-doped iron oxide nanoparticles had a lattice parameter of 8.3830 Å and a crystallite size of 14.1 nm, Wustite XRD pattern had a crystallite size of 15.2 nm with a lattice parameter of 4.1867 Å, and iron oxide nanoclusters showed a crystallite size of 6.5 nm and a lattice parameter of 8.3432 Å. Transmission electron microscopy images shown below for each particle type with 50 nm scale bars.

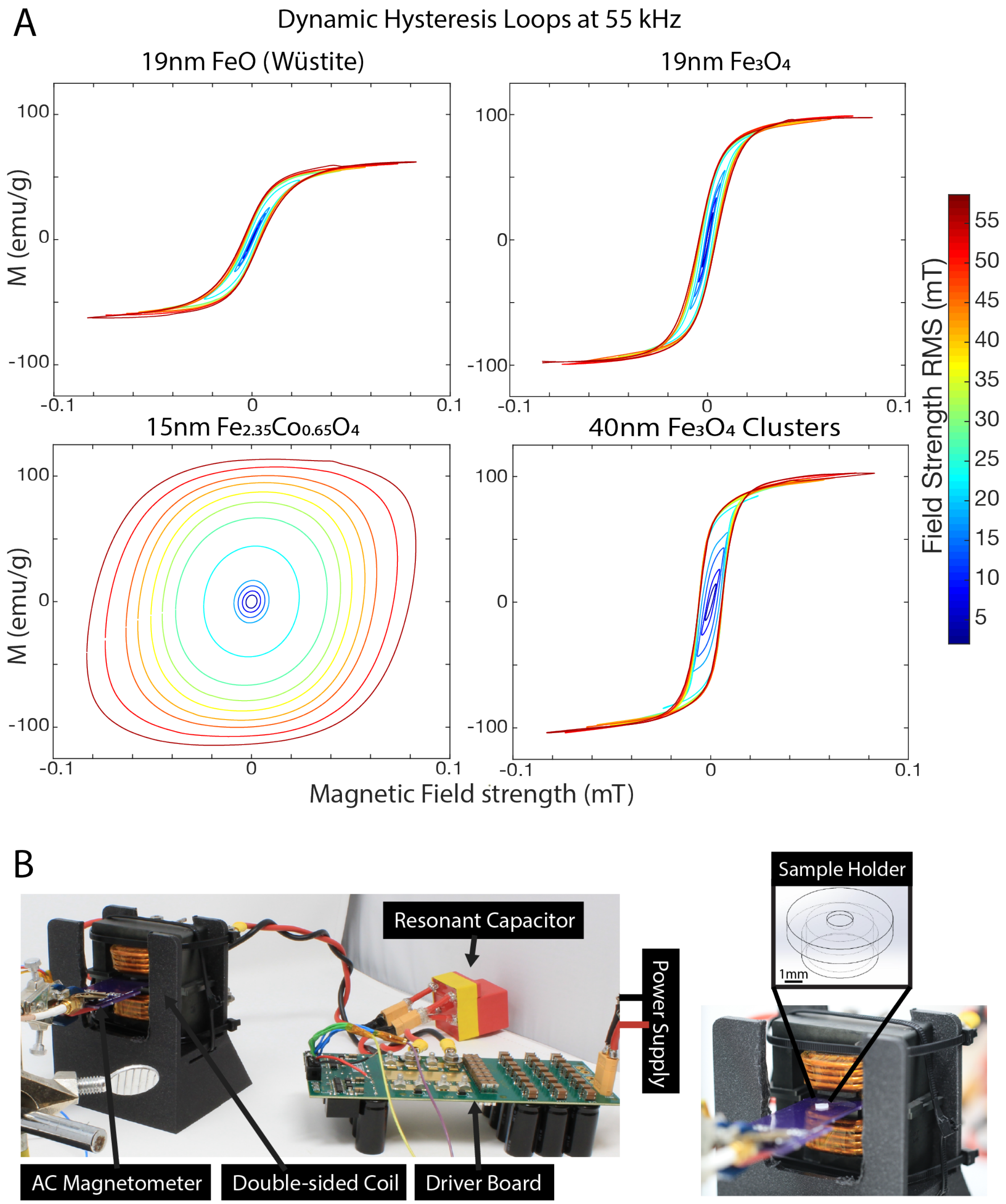


**Supplemental Figure S5 | Dynamic hysteresis loops of nanoparticles measured by AC Magnetometer in AMF at 55 kHz**

**A)** Room temperature (22-25 °C) analysis of 19nm FeO and Fe3O4, 15nm Cobalt-doped iron oxide, and 40nm Fe3O4 nanoclusters showing hysteresis loops measured from 17 μL of each sample recorded at ~10 mg/mL as measured by custom AC magnetometer set up. Hysteresis loops show clear coercivity differences compared to iron oxide nanoparticles that saturate at lower magnetic field strengths. **B)** AC magnetometer set up showing AMF field generation with custom driver, double-sided coil around N87 ferrite cores, resonant capacitor, driver boards, power supply leads, and 3D printed sample holder.

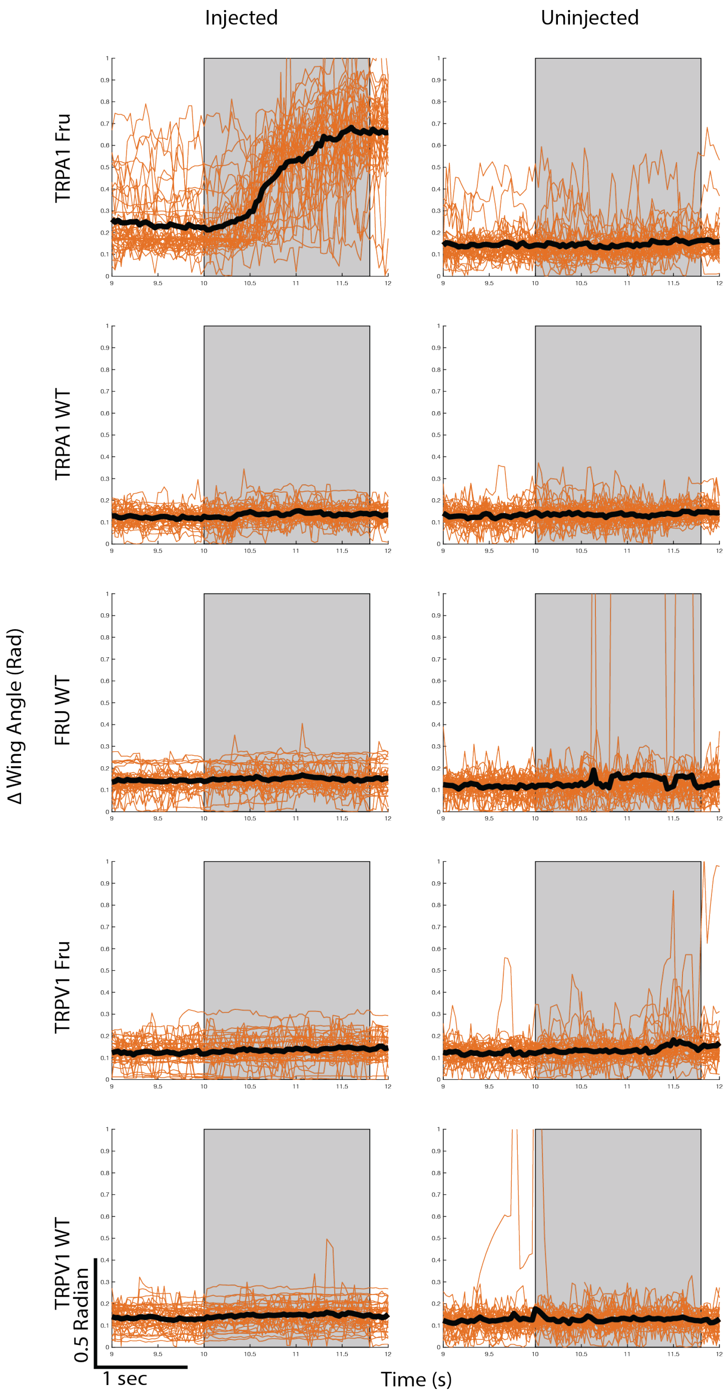


**Supplemental Figure S6 | DLC Wing Angle Traces of Experimental vs Control Flies.**
Average Δ wing angle measurements (black trace) from 10 flies for each condition, quantified using DeepLabCut. The Δ wing angle trace for each AMF stimulation of each fly is represented in orange and the AMF stimuli is indicated by the grey box. Each fly received 4 AMF stimulations of 1.8 s (49.9 kHz; 80 mT). No significant wing opening response during AMF is observed for any of the control groups.

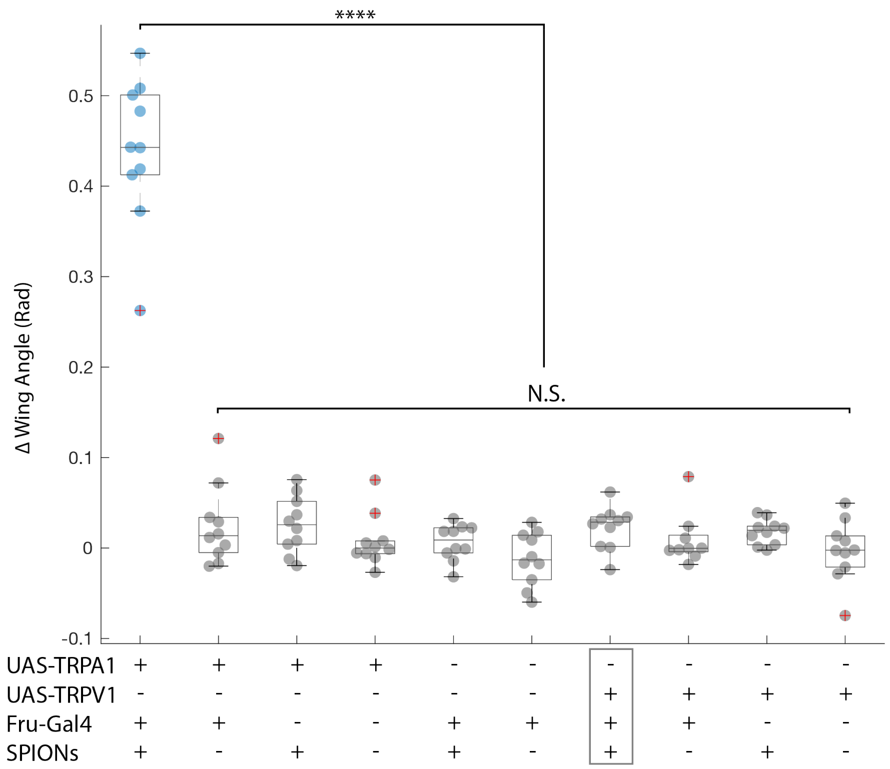


**Supplemental Figure S7 | Statistical Comparison of Wing Angle Opening During AMF Stimulation**
Average Δ wing angle measurements of 10 flies for each group. Each fly is represented by the average of 4 AMF stimulations of 1.8 s (49.9 kHz; 80 mT) and is quantified using DeepLabCut. No significant wing opening response during AMF is observed for any of the control groups, including the group injected with SPIONs and human TRPV1 under the fruitless driver (grey box). One-way ANOVA of delta wing angle taken immediately before and after stimulation and compared to uninjected control flies (**** = $p < 0.0001$). Normal distribution for each group determined by Chi-square goodness-of-fit test. Post-hoc analysis with Tukey HSD Test. Outliers are marked with red + if greater than [q3 + 1.5*(q3-q1)] or smaller than [q1 - 1.5*(q3-q1)] but still included in statistical analysis.

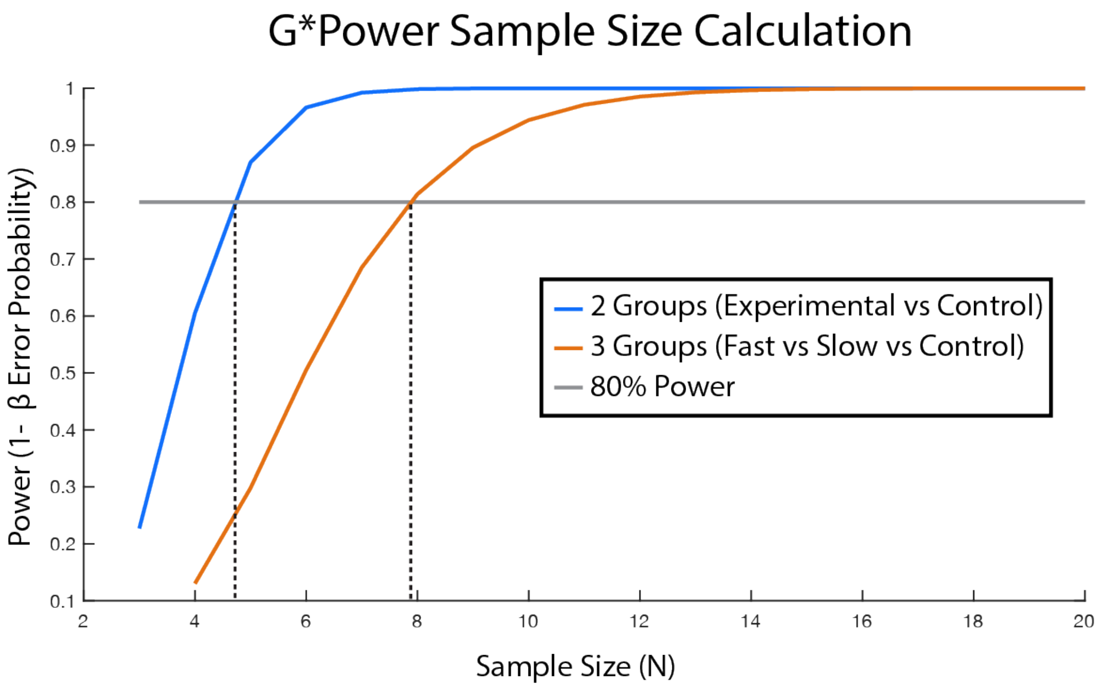


**Supplemental Figure S8 | Sample Size Calculation using G*Power**
Sample sizes were calculated using G*Power using preliminary data from 5 flies in each group (Control vs Experimental). Sample sizes were calculated using the 'ANOVA: Fixed effects, omnibus, one-way' test with an alpha error probability of 0.05 and power as 0.8. The 2-group test was conducted with just the fast ramp data vs uninjected controls resulting in an effect size of f=2.12. The 3-group test was conducted comparing fast and slow ramp data versus the uninjected controls resulting in an effect size of f=1.56. This concluded that sample sizes of 5+ would work for experimental vs control and sample sizes of 8+ would be sufficient to compare slow vs fast ramps and controls. We exceeded this and used 10 flies to compare against control and 20 experimental flies to compare between the different thermal ramp conditions.

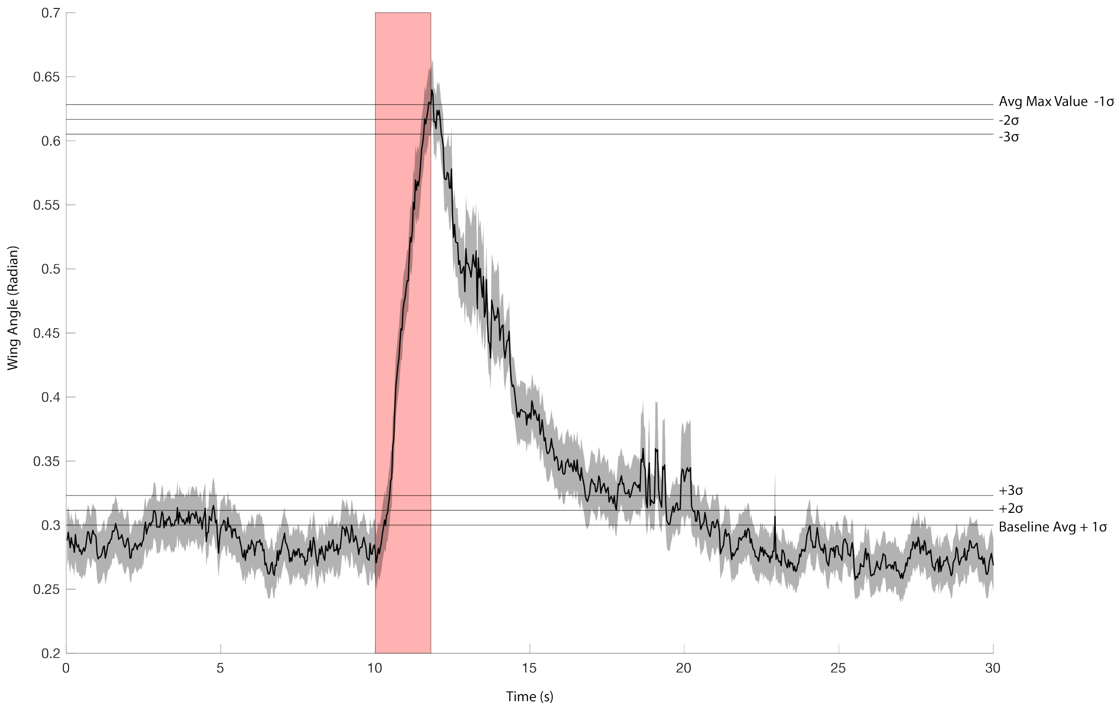


| | On Time | Off Time (Max - σ) | Return to baseline (Avg + δ) |
|---|---|---|---|
| **1 Sigma (68%)** | **~300ms** | **100ms** | **8.73s** |
| **2 Sigma (95%)** | **~430ms** | **100ms** | **7.63s** |
| **3 Sigma (99.7%)** | **~500ms** | **370ms** | **5.00s** |

**Supplemental Figure S9 | On/Off Temporal Latency of Average Wing Opening Response**
Average wing angle measurements from magnetically stimulated drosophila expressing TRPA1 under the fruitless driver and injected with cobalt doped iron oxide nanoparticles. The wing angle changes are quantified using DeepLabCut on 40 experiments consisting of 2x AMF stimulations of 1.8 s (49.9 kHz; 80 mT). The "On Time" is reported here as the first time the average trace crosses the baseline average + 1,2, or 3σ. The σ value is calculated as the wing angle standard deviation during the 10 seconds before each AMF stimulation. The "Off Time" specifies when the average plot drops 1,2, or 3 σ below the average max value and the "Return to baseline" indicates the first time the average trace drop below average baseline + 1, 2, or 3σ. The greyed-out area indicates the standard error of the mean.

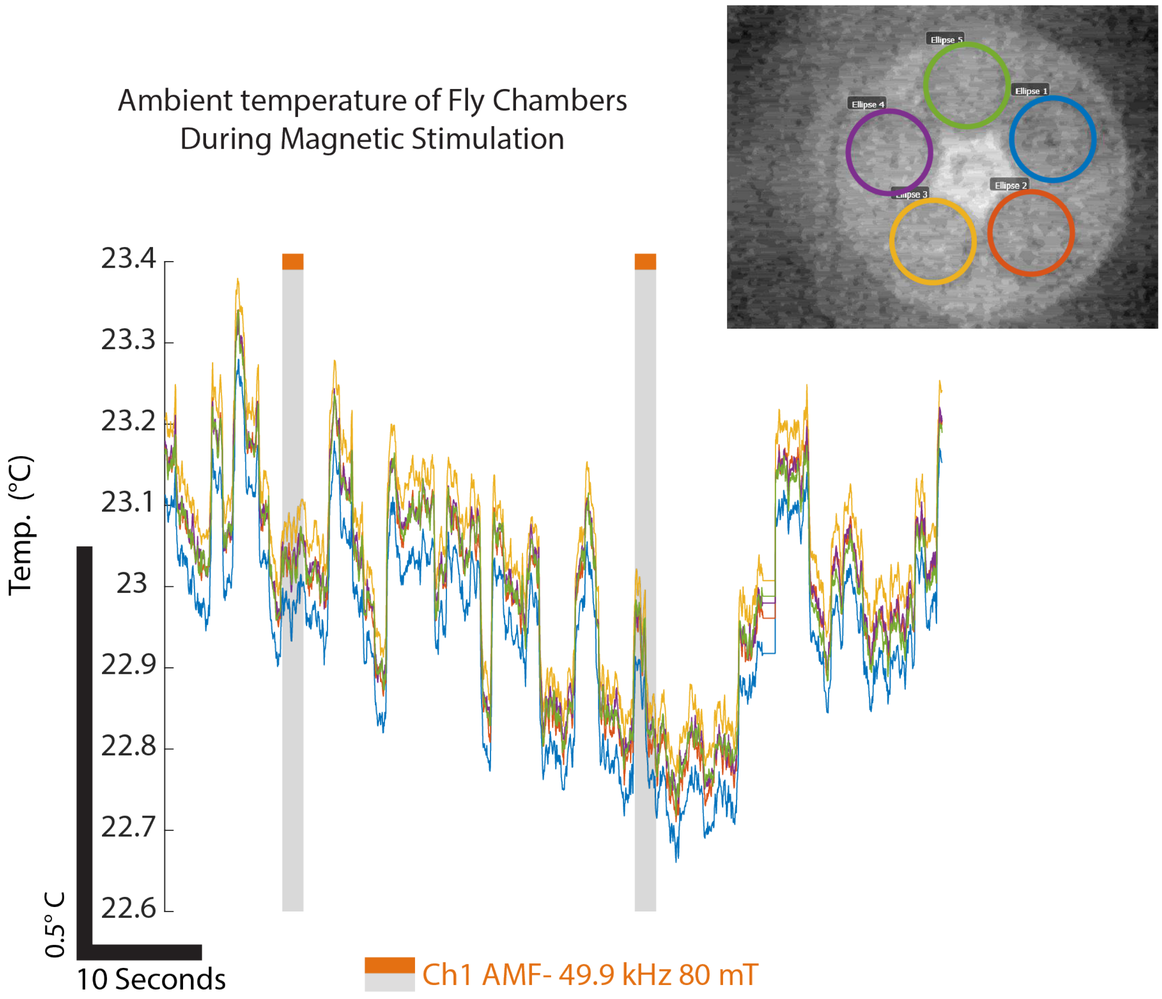


**Supplemental Figure S10 | Infrared Thermal Imaging of Fly Chambers During Magnetic Stimulation**
Thermal imaging traces of each chamber shown recorded with infrared fluorescent camera (FLIR A700). Noise in the sensor show fluctuations of about 0.5 °C and there is not noticeable difference in temperature of each chamber during magnetic stimulation above the noise floor. The acrylic cap above the chamber was removed to record from the bottom layer of the chamber that sits just below the fly during recording.

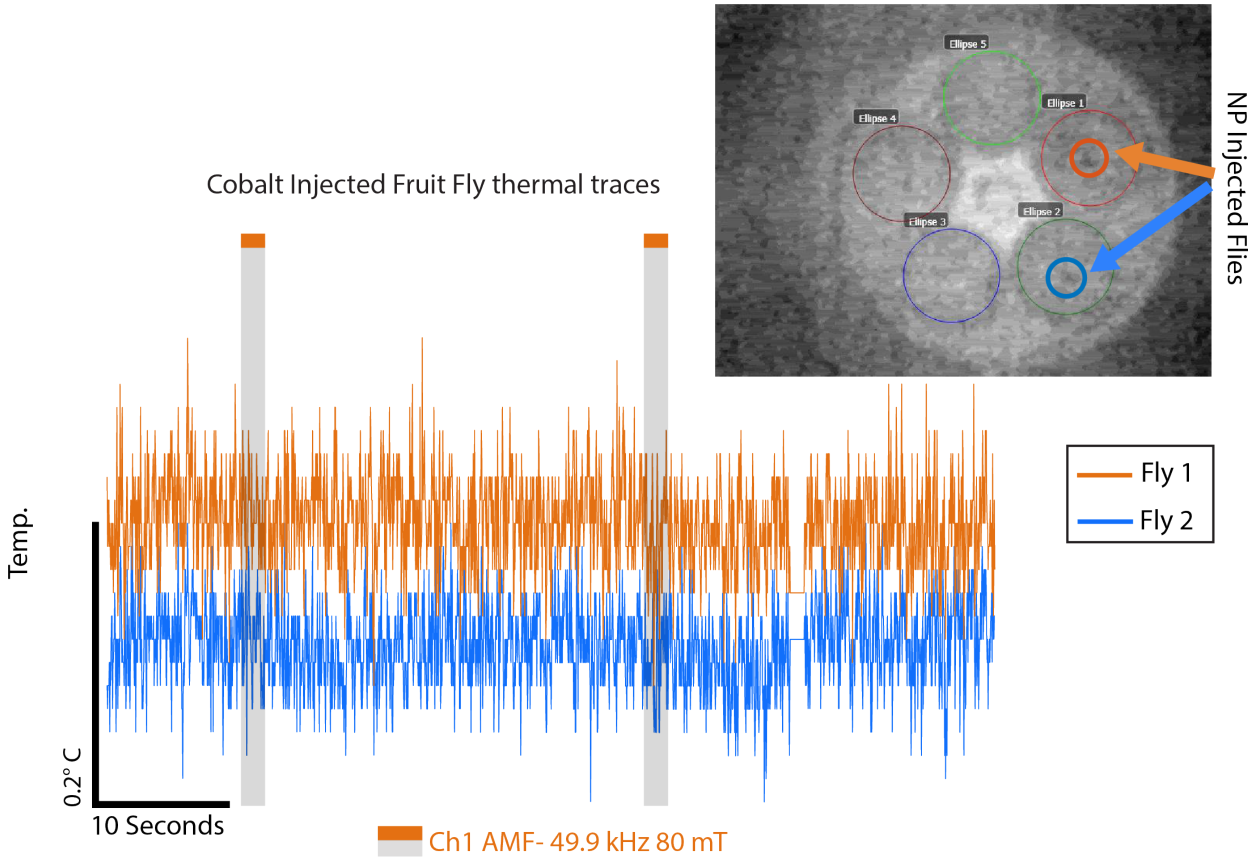


**Supplemental Figure S11 | Infrared Thermal Imaging of Injected *Drosophila* During Magnetic Stimulation**
Average thermal imaging traces of injected drosophila shown after subtracting background noise using an ROI elsewhere within the chamber. Previously responsive *Drosophila* injected with nanoparticles were immobilized before being placed in the chamber and show no noticeable heating during AMF stimulation above the noise of the sensor (FLIR A700).

**Supplemental Video S1 | Slow Thermal Ramp of Cobalt Injected *Drosophila* for *Fru* circuit stimulation**
**Left:** Shows chamber containing 5 flies expressing TRPA1-A channels from the UAS promoter driven by *Fru-GAL4*. Stimulation of this circuit results in a wing extension. 15nm Cobalt-doped iron oxide nanoparticles ($Fe_{2.35}Co_{0.65}O_4$) injected flies are shown within orange circles. **Right:** DeepLabCut analysis of fruit fly wing extension (θ) shows traces of wing extensions when exposed to 20 second pulses of AMF stimulation (19 mT; 49.9 kHz) shown in orange boxes. The responses are muted because slow ramp stimulation does not efficiently activate the rate sensing TRPA1-channel. The white scrub line shows where the video lines up with the traces. Top to bottom traces correspond to chambers top 3 chambers left to right followed by the bottom 2 chambers left to right.

**Supplemental Video S2 | Fast Thermal Ramp of Cobalt Injected and Uninjected *Drosophila* for *Fru* circuit stimulation**
**Left:** Top chamber contains the same 5 flies as in Supplemental Video S1, expressing TRPA1-A channels from the UAS promoter driven by *Fru-GAL4*. Stimulation of this circuit results in a wing extension. 15nm Cobalt-doped iron oxide nanoparticles ($Fe_{2.35}Co_{0.65}O_4$) injected flies are shown within orange circles at the top. Bottom chamber shows 5 uninjected control flies expressing TRPA1-A driven by a *Fru-Gal4* promoter shown within grey circles at the bottom. **Right:** DeepLabCut analysis of fruit fly wing extension (θ) shows traces of wing extensions when exposed to 1.8 second pulses of AMF stimulation (80 mT; 49.9 kHz) shown in orange boxes. The white scrub line shows where the video lines up with the traces. Top to bottom traces correspond to chambers top 3 chambers left to right followed by the bottom 2 chambers left to right for each chamber of flies. Video is shown at 2x speed.

**Supplemental Video S3 | *Drosophila Hb9* circuit stimulation with wüstite injected controls**
Shows chamber containing 5 flies expressing TRPA1-A channels from the UAS promoter driven by *Hb9-GAL4*. Stimulation of this circuit results in side-to-side movement. 19nm magnetite ($Fe_3O_4$) injected flies are shown within blue circles. 19nm wüstite (FeO – non-heating control nanoparticles) injected flies are shown within black circles. Positions of the flies were tracked with FlyTracker and overlaid on top of the flies showing the previous 10 seconds of fly movement during the video. Flies are exposed to a single 30 second pulse of the alternating magnetic field (40 kA/m or ~50mT; 380 kHz).

**Supplemental Video S4 | Multiplexed *Drosophila Fru* circuit stimulation**
**Left:** Shows chamber containing 5 flies expressing TRPA1-A channels from the UAS promoter driven by *Fru-GAL4*. Stimulation of this circuit results in a wing extension. 15nm Cobalt-doped iron oxide nanoparticles ($Fe_{2.35}Co_{0.65}O_4$) injected flies are shown within orange circles and 40nm iron oxide nanocluster ($Fe_3O_4$) injected flies are shown in blue circles. **Right:** DeepLabCut analysis of fruit fly wing extension (θ) shows traces of wing extensions when exposed to 3 second pulses of AMF stimulation for channel 1 (80 mT; 49.9 kHz) shown in orange boxes and stimulation for channel 2 (12 mT; 555 kHz) shown in blue boxes. The white scrub line shows where the video lines up with the traces. Top to bottom orange traces correspond to chambers top 3 chambers left to right followed by the bottom 2 chambers left to right.
